# Astrocytes Contribute to Remote Memory Formation by Modulating Hippocampal-Cortical Communication During Learning

**DOI:** 10.1101/682344

**Authors:** Adi Kol, Adar Adamsky, Maya Groysman, Tirzah Kreisel, Michael London, Inbal Goshen

**Affiliations:** Edmond and Lily Safra Center for Brain Sciences (ELSC), The Hebrew University of Jerusalem, Jerusalem, 91904 Israel; ELSC Vector Core Facility, The Hebrew University of Jerusalem, Jerusalem, 91904 Israel; Alexander Silberman Institute of Life Sciences, The Hebrew University of Jerusalem, Jerusalem, 91904 Israel

**Keywords:** Astrocytes, Hippocampus, Anterior Cingulate Cortex (ACC), Fear conditioning, Remote Memory, Non Associative Place Recognition, In-Vivo Recording, Chemogenetics, hM4Di, Optogenetics, cFos, Neurogenesis.

## Abstract

The consolidation and retrieval of remote memories depend on the coordinated activity of the hippocampus and frontal cortices. However, the exact time at which these regions are recruited to support memory and the interactions between them are still debated. Astrocytes can sense and modify neuronal activity with great precision, but their role in cognitive function has not been extensively explored. To investigate the role of astrocytes in remote memory we expressed the Gi-coupled receptor hM4Di in CA1 astrocytes, allowing their manipulation by a designer drug. We discovered that astrocytic modulation during learning resulted in a specific impairment in remote, but not recent, memory recall, accompanied by decreased neuronal activity in the anterior cingulate cortex (ACC) during retrieval. We revealed a massive recruitment of ACC-projecting neurons in CA1 during memory acquisition, accompanied by activation of ACC neurons. Astrocytic Gi activation disrupted CA3 to CA1 communication in-vivo, and reduced the downstream response in the ACC. This same manipulation in behaving mice induced a projection-specific inhibition of ACC-projecting CA1 neurons during learning, consequently preventing the recruitment of the ACC. Our findings suggest that the foundation of remote memory is established in the ACC during acquisition, engaging a distinct process from the one supporting consolidation of recent memory. Furthermore, the mechanism underlying remote memory involves projection-specific functions of astrocytes in regulating neuronal activity.

## INTRODUCTION

Remote memories, weeks to decades long, continuously guide our behavior, and are critically important to any organism, as the longevity of a memory is tightly connected to its significance. The ongoing interaction between the hippocampus and frontal cortical regions has been repeatedly shown to transform in the transition from recent (days long) to remote memory^1–3^. However, the exact time at which each region is recruited, the duration for which it remains relevant to memory function, and the interactions between these regions, are still debated.

Astrocytes are no longer considered to merely provide homeostatic support to neurons and encapsulate synapses, as pioneering research has shown that astrocytes can also sense and modify synaptic activity as an integral part of the ‘tripartite synapse’^4, 5^. Interestingly, astrocytes demonstrate extraordinary specificity in their effects on neuronal circuits^6^, at several levels: First, astrocytes differentially affect neurons based on their genetic identity. For example, astrocytes in the dorsal striatum selectively respond to, and modulate, the input onto two populations of medium spiny neurons, expressing either D1 or D2 dopamine receptors^7^. Similarly, astrocytes selectively modulate the effects of specific inhibitory cell-types, but not others, in the same brain region^8–11^. Second, astrocytes exert neurotransmitter-specific effects on neuronal circuits. For instance, astrocytic activation in the central amygdala specifically depresses excitatory inputs and enhances inhibitory inputs. Finally, astrocytes exhibit task-specific effects in-vivo, i.e. astrocytic stimulation selectively increases neuronal activity when coupled with memory acquisition, but not in the absence of learning^12^. An intriguing open question is whether astrocytes can differentially affect neurons based on their distant projection target.

The integration of novel chemogenetic and optogenetic tools in astrocyte research allows real-time, reversible manipulation of these cells at the population level, in combination with electrophysiological and behavioral measurements. Such tools were used in brain slices to activate intracellular pathways in astrocytes, and show the ability of these cells to selectively modulate the activity of the neighboring neurons in the amygdala and striatum^13, 14^, and induce de-novo long-term potentiation in the hippocampus^12, 15^. The reversibility of chemogenetic and optogenetic tools allows careful dissection of the effect of astrocytes during the different stages of memory in behaving animals^16, 17^. The recruitment of different intracellular signaling pathways in astrocytes using such tools is starting to shed light on their complex involvement in memory processes, with Gq activation in the CA1 during acquisition (but not during recall) resulting in enhanced recent memory^12, 15^, and Gs activation resulting in recent memory impairment^18^.

To explore the role of astrocytes in remote memory, and their ability to exert projection-specific effects, we used chemogenetics to activate the Gi pathway in these cells, and found that this astrocytic modulation in CA1 during learning resulted in a specific impairment in remote (but not recent) memory recall, accompanied by decreased activity in the anterior cingulate cortex (ACC) at the time of retrieval. In-vivo Gi activation in astrocytes disrupted synaptic transmission from CA3 to CA1 and reduced the downstream recruitment of the ACC. Finally, we show a dramatic recruitment of CA1 neurons projecting to ACC during memory acquisition, and a projection-specific inhibition of this population by Gi pathway activation in CA1 astrocytes.

## RESULTS

### Gi pathway activation in CA1 astrocytes specifically impairs the acquisition of remote memory

To specifically modulate the activity of CA1 astrocytes via the Gi pathway we employed an AAV8 vector encoding the designer receptor hM4Di fused to mCherry under the control of the astrocytic GFAP promoter. Stereotactic delivery of this AAV8-GFAP::hM4Di-mCherry vector resulted in CA1-specific expression restricted to astrocytic outer membranes (Fig. 1A,B), with high penetrance (>85% of the GFAP cells expressed hM4Di; Fig. S1A), and the promoter provided almost complete specificity (>95% hM4Di positive cells were also GFAP positive; Fig. S1B). Co-staining with the neuronal nuclei marker NeuN showed no overlap with hM4Di expression (Figure S1C).

**Figure 1.**
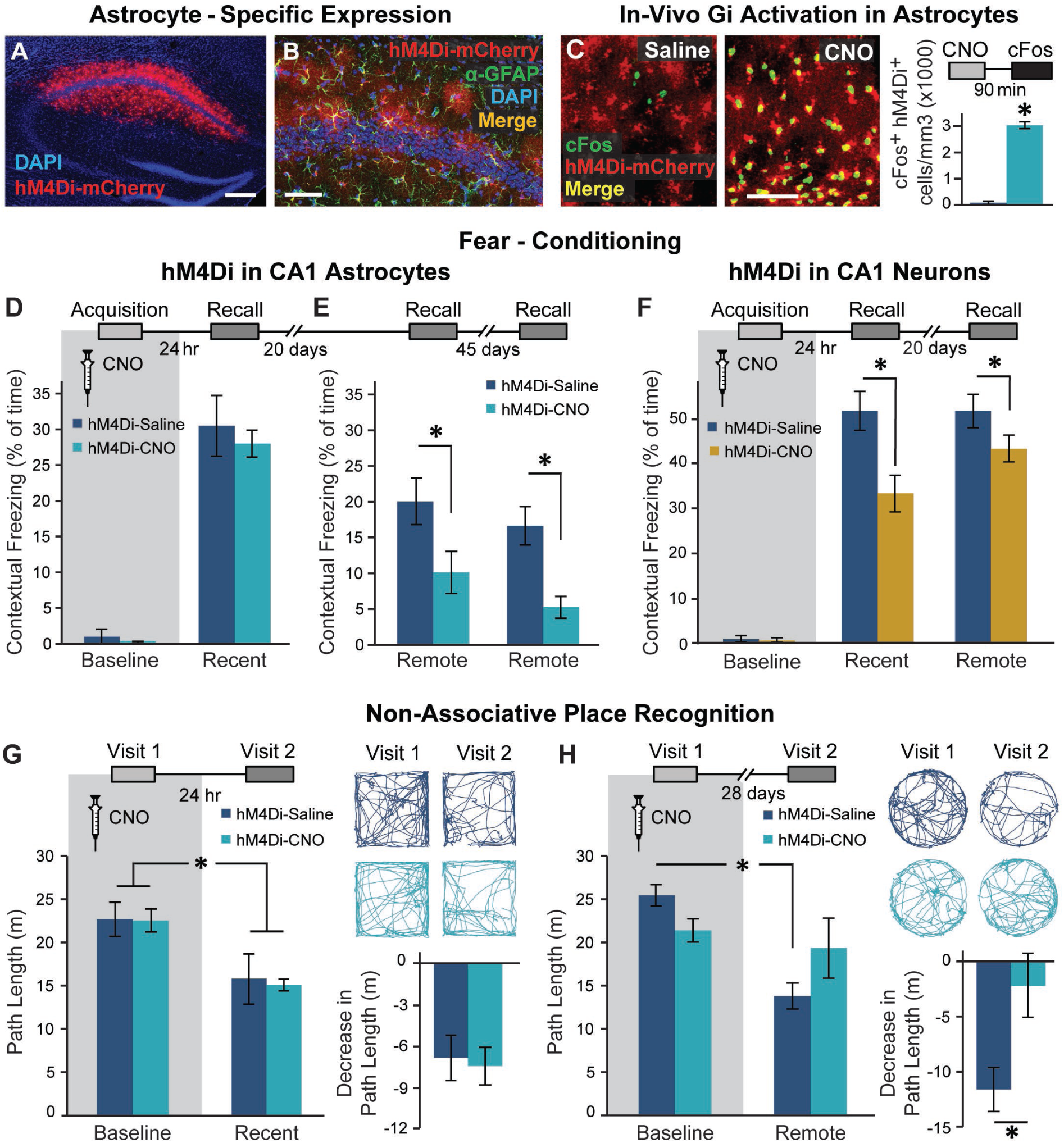
Astrocytic Gi pathway activation in CA1 during learning specifically impaired remote contextual memory. (**A**) Bilateral double injection of AAV8-GFAP::hM4Di-mCherry resulted in hM4Di expression selectively in CA1 (scale bar 200µm). (**B**) hM4Di (red) was expressed in the astrocytic membrane around the soma, as well as in the distal processes (scale bar 50µm). (**C**) CNO administration in-vivo to mice expressing hM4Di (red) in CA1 astrocytes resulted in a significant increase in cFos expression (green) in these astrocytes, compared to saline injected controls (p<0.00005, n = 2-4 mice, 6-15 slices per groups; scale bar 50μm). (**D**) Mice expressing hM4Di in their CA1 astrocytes were injected with either Saline (n=6) or CNO (n=6) 30min before fear conditioning (FC) acquisition. CNO application before training had no effect on baseline freezing before shock administration or on recent contextual freezing on the next day compared to Saline treated controls. (**E**) CNO application before training resulted in a >50% impairment (p<0.05) in contextual freezing in CNO-treated mice tested 20 days later, compared to Saline treated controls (*left*). An even bigger impairment of >68% (p<0.005) was observed 45 days later (*right*). (**F**) Mice expressing hM4Di in their CA1 neurons were injected with either Saline (n=9) or CNO (n=10) 30min before FC acquisition. CNO application before training had no effect on baseline freezing before shock administration, bur resulted in decreased recent contextual freezing on the next day (p<0.005), and decreased remote recall 20 days after that (p<0.05) compared to Saline treated controls. (**G**) In the non-associative place recognition test, astrocytic Gi pathway activation by CNO application before a first visit to a new environment had no effect on recent memory, reflected by a similar decrease (p<0.0001) in the exploration between Saline injected (n=6) and CNO-treated mice (n=8). Example exploration traces and the average change (Δ) in exploration following treatment are shown on the right. (**H**) Astrocytic modulation impaired remote recognition of the environment on the second visit, reflected by a decrease in the exploration only in the Saline injected (n=6)(p<0.01), but not CNO-treated (n=6) mice. Example exploration traces and average decrease Δ are shown on the right. Data presented as mean ± standard error of the mean (SEM).

Recent work has shown that hM4Di activation in astrocytes mimics the response of these cells to GABAergic stimuli^14, 19^, and induces elevated expression of the immediate-early gene cFos in-vivo^14, 19, 20^. To verify this effect in our hands, mice were injected with CNO (10mg/kg, i.p.), brains were collected 90 min later and stained for cFos. As expected, CNO dramatically increased cFos levels in astrocytes of hM4Di-expressing mice, compared to saline-injected controls (Figure 1C; p<0.00005, t-test). As cFos is similarly induced by the recruitment of the Gq pathway^12, 20^, it seems not to be a reliable indicator of the nature of astrocytic activity manipulation, but only to the occurrence of a significant modulation.

Previous elegant research demonstrated the necessity of normal astrocytic metabolic support to memory and showed that chronic genetic manipulations in astrocytes can affect recent memory^21–27^. The contribution of astrocytes to remote memory, however, was never investigated. To address this topic, we took advantage of the temporal flexibility offered by chemogenetic tools, allowing not only cell-specific, but also memory-stage specific (e.g. during acquisition or recall), reversible modulation of astrocytes^12, 13^.

To test the effect of astrocytic modulation on cognitive performance, mice were injected bilaterally with AAV8-GFAP::hM4Di-mCherry into the dorsal CA1, and three weeks later CNO (10mg/kg, i.p.) was administered 30 minutes before fear conditioning (FC) training, in which a foot-shock was paired with a novel context and an auditory cue. CNO application in GFAP::hM4Di mice had no effect on the exploration of the conditioning cage before shock administration (Figure S1D) or on baseline freezing before shock delivery (Figure 1D *left*).

One day later, when CNO was no longer present^28, 29^, mice were placed back in the conditioning context and freezing was quantified. We found no difference in recent memory retrieval between GFAP::hM4Di mice treated with CNO or with saline during FC acquisition (Figure 1D *right*). Remarkably, when the same mice were tested in the same context 20 days later, those treated with CNO during conditioning showed a dramatic impairment in memory retrieval (Figure 1E *left*; p<0.05, t-test). This deficiency was still clearly observed 45 days after that, when these mice were re-tested in the same context for a third time (Figure 1E *right*; p<0.005, t-test). The effect of CA1 astrocytic manipulation was unique to the hippocampal-dependent contextual memory task, as no effect was observed when the same mice were tested for auditory-cued memory in a novel context, i.e. both groups demonstrated similar freezing in response to the tone one day after training (Figure S1E; F_(1,10)_=79.84, time main effect, p<0.001), and 20 days later (Figure S1F; F_(1,10)_=10.00, time main effect, p<0.01).

We then tested what effects would inhibition of CA1 neurons have on recent and remote memory recall. We injected mice with an AAV5-CaMKIIα::hM4Di-mCherry vector to induce hM4Di expression in CA1 glutamatergic neurons (Figure S1G). To test the effect of direct neuronal inhibition on recent and remote memory acquisition, we injected CaMKIIα::hM4Di mice with CNO (10mg/kg, i.p.) 30 minutes before FC acquisition. Gi pathway activation in neurons had no effect on the exploration of the conditioning cage before tone and shock administration (Figure S1H), or on baseline freezing levels (Figure 1F *left*). Mice were then fear-conditioned, and tested on the next day. As expected, neuronal inhibition during training resulted in impaired contextual freezing one day later (Figure 1F *middle*; p<0.005, t-test). When the same mice were tested in the same context 20 days later, the memory impairment was still apparent (Figure 1F *right*; p<0.05, t-test). No significant effect on auditory-cued memory in a novel context was observed, at either the recent or the remote time points, as both groups demonstrated similar freezing in response to the tone (Figure S1I-J; F_(1,17)_=155.44, time main effect p<0.000001, F_(1,17)_=34.72, time main effect p<0.00001, respectively). Thus, general neuronal inhibition during acquisition impairs both recent and remote memory.

Effects specific to remote, but not recent, memory were reported in the past in response to neuronal manipulations during recall (e.g. ^30–32^). Based on these reports, we next tested the necessity of intact astrocytic function during the retrieval of recent and remote memory, by administering CNO during the recall tests. CNO administration during recent and remote recall of contextual and auditory-cued memory had no effect on freezing levels compared to saline-injected controls (Figure S1K-M). Thus, normal astrocytic activity is not required during either recent or remote memory recall, but only during memory acquisition.

To further validate the unexpected effect of astrocytic Gi pathway activation during acquisition on remote memory in a less stressful task, we employed an additional paradigm, the ’non-associative place recognition’ (NAPR) test. In this task, mice are allowed to explore a novel open field, and upon re-exposure to the same arena are expected to display decreased exploration of this now familiar environment. Indeed, GFAP::hM4Di mice injected with either saline or CNO during NAPR acquisition showed a marked decrease in exploration upon a second exposure to the square environment to which they were exposed 24 hours earlier, as expected (Figure 1G; F_(1,12)_=45.69, no interaction, time main effect p<0.0001). Another cohort of GFAP::hM4Di mice injected with saline during NAPR acquisition showed a marked decrease in exploration upon the second exposure to a round environment to which they were introduced 4 weeks earlier, as expected. However, exploration level in GFAP::hM4Di mice treated with CNO did not decrease (Figure 1H *left*), suggesting that they did not recall the remote original experience in this context. These findings were reflected in a significant treatment by time interaction (F_(1,10)_=5.890, p<0.05), and post-hoc analysis showed a significant difference between the first and second visit only for the saline group (p<0.01). A significant effect was also found for the decrease in exploration of saline and CNO treated mice (p<0.01, t-test; Figure 1H *right*). To confirm that these mice are still capable of performing the NAPR task normally when astrocytic activity is intact, and verify the absence of non-specific long-term effects, we repeated the experiment in a novel trapezoid environment with no CNO administration during the first visit, in the same cohort, which now demonstrated comparable performance between groups (Figure S1N; F_(1,10)_=11.855, time main effect p<0.01, no interaction).

To verify that our results did not stem from the CNO application itself, control mice injected with an AAV8-GFAP::eGFP vector (Figure S2A) were trained in the same behavioral paradigms. CNO administration (10mg/kg, i.p.) in these GFAP::eGFP mice had no effect on baseline freezing, recent or remote contextual memory (Figure S2B-C), or on performance in the NAPR task upon their second visit to this environment a month later (Figure S2D; F_(1,11)_=58.66, time main effect p<0.0001, no interaction).

Our results show that Gi activation in CA1 astrocytes during the acquisition of spatial memory selectively impairs its remote, but not recent, recall, whereas direct neuronal inhibition during acquisition impairs both recent and remote memory. These findings raise two novel hypotheses: First, that the foundation for remote memory is established during acquisition, in a parallel separate process to recent memory, and can thus be manipulated independently. And second, that astrocytes are able to specifically modulate the acquisition of remote memory, with precision not granted by general neuronal inhibition. Both hypotheses are tested below.

### Astrocytic Gi pathway activation during memory acquisition reduces the recruitment of brain regions involved in remote memory, during retrieval

The transition from recent to remote memory is accompanied by brain-wide reorganization, including the recruitment of frontal cortical regions like the ACC^1–3, 31, 33, 34^, indicated by increased expression of cFos^31, 33^. To gain insight into changes in the neuronal activity accompanying the recent and remote retrieval of memories acquired under astrocytic modulation, GFAP::hM4Di mice were injected with Saline or CNO before FC acquisition, brains were collected 90 minutes after recent or remote recall, and stained for cFos (Figure 2B *top*). We quantified retrieval-induced cFos expression in neurons at CA1 and ACC (Figure 2A), area repeatedly implicated in remote memory^2, 35^. As before, CNO administration to GFAP::hM4Di mice during acquisition had no effect on recent contextual memory (Figure 2B *bottom*), and changes in cFos expression following recent recall in either CA1 or ACC were not observed (Figure 2C-E). Another cohort of GFAP::hM4Di mice was injected with CNO before acquisition, tested for recent memory 24 hours later, and then for remote recall 21 days after that. Importantly, we replicated our initial finding that astrocytic modulation during acquisition specifically impaired remote but not recent contextual memory (Figure 2F; p<0.05, t-test). Impaired remote memory was accompanied by reduced cFos expression in both the CA1 (p<0.05, t-test) and the ACC (p<0.01, t-test) regions (Figure 2G-I). We also performed the same cFos quantification in brains collected after the last recall test from the first behavioral experiment (Figure 1E), of mice that were injected with CNO >60 days earlier. In this experiment too, impaired remote recall in GFAP::hM4Di mice treated with CNO during conditioning was accompanied by reduced cFos expression in CA1 and ACC compared to saline treated mice (Figure S3B; p<0.05 for both, t-test).

**Figure 2.**
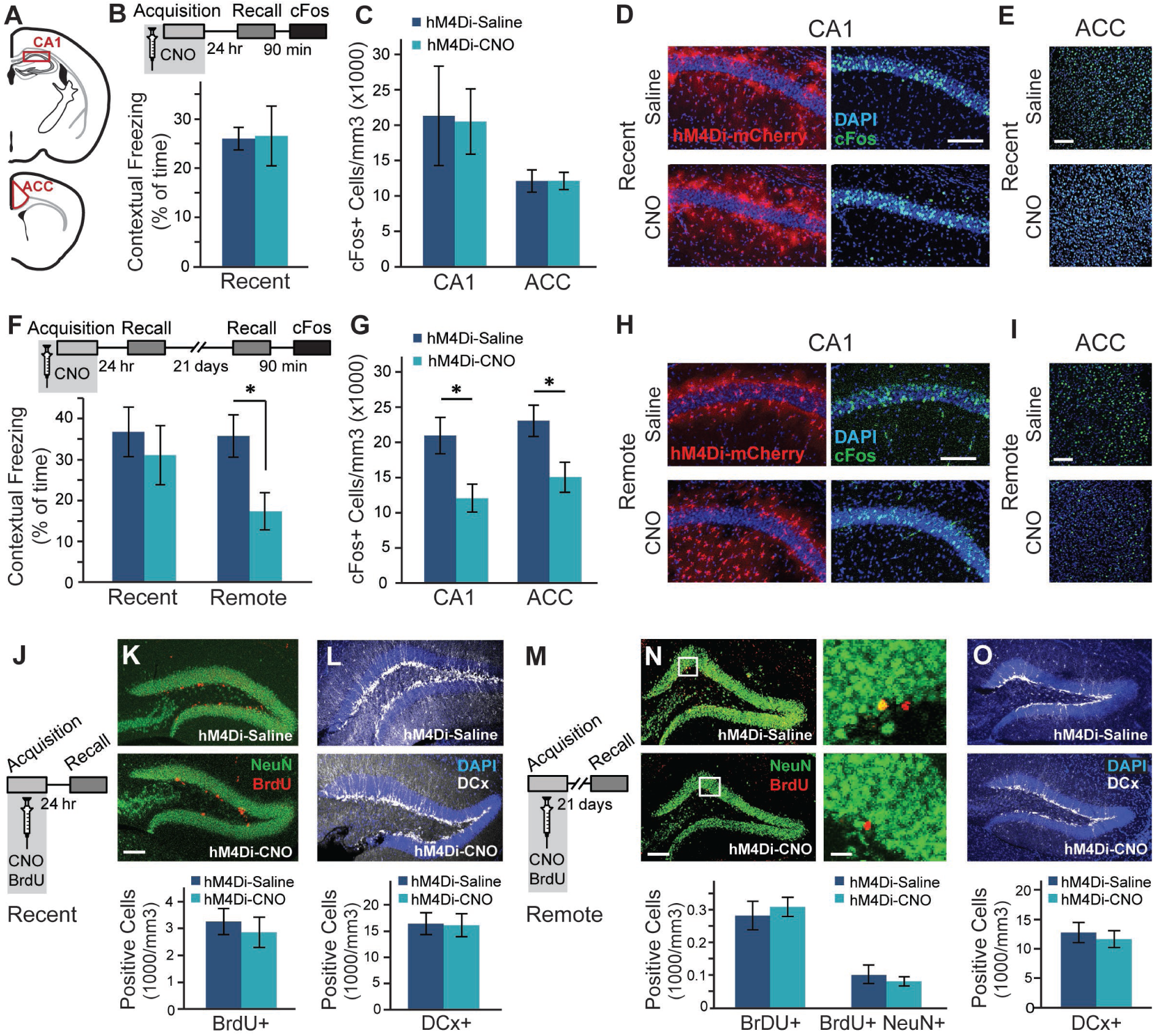
Astrocytic Gi activation during memory acquisition reduced CA1 and ACC activity at the time of remote recall, but did not affect neurogenesis. (**A**) Active neurons expressing cFos were quantified in the CA1 and ACC regions. GFAP::hM4Di mice were injected with CNO (n=5) or Saline (n=5) before fear conditioning, and then tested on the next day. No changes were observed in recent memory (**B**) or in the number of neurons active during recall in the CA1 or ACC (**C**). Representative images of hM4Di (red) and cFos (green) in the CA1 (**D**) and ACC (**E**) are presented. Other GFAP::hM4Di mice were injected with CNO (n=5) or Saline (n=6) before fear conditioning, and then tested on the next day and again 21 days later. No changes were observed in recent memory (F *left*). However, CNO application before training resulted in >50% reduction (p<0.05) in contextual freezing 21 days later, compared to Saline treated controls (F *right*). Impaired remote recall was accompanied by reduced number of cFos-expressing neurons in CA1 and ACC (p<0.05 and p<0.01, respectively)(**G**). Representative images of the CA1 (**H**) and ACC (**I**) are presented. (**J**) GFAP::hM4Di mice were injected with CNO (n=5) or Saline (n=5) together with BrdU before fear conditioning, and then tested on the next day. No changes were observed in stem cell proliferation (Brdu in red)(**K**) or in the number of young, Doublecortine (DCx)-positive neurons (white)(**L**). (**M**) GFAP::hM4Di mice were injected with CNO (n=5) or Saline (n=6) and BrdU before fear conditioning, and then tested 21 days later. No changes were observed in stem cell proliferation and differentiation (**N**) or in the number of young, DCx-positive neurons (**O**). All scale bars = 100μm, except zoomed-in image in panel N where scale bar = 10μm. Data presented as mean ± SEM.

In the same mice we also quantified retrieval-induced cFos expression in several additional brain regions known to be involved in memory: the Dentate Gyrus (DG) of the hippocampus, the Retrosplenial Cortex (RSC), and the Basolateral Amygdala (BLA) (Figure S3A). No changes in cFos expression in the DG or RSC were observed (Figure S3C). BLA cFos expression was reduced in GFAP::hM4Di mice treated with CNO (p<0.05, t-test; Figure S3C), which may be attributed to the reduced fear. Finally, to exclude any non-specific effects of CNO itself, we repeated the same experiments in control GFAP::eGFP mice. As before, CNO application induced no difference in either recent or remote fear memory, and we found no alterations in cFos expression (Figure S3D-K).

Again, we show that astrocytic Gi pathway activation during fear memory acquisition selectively impaired remote recall, but spared recent retrieval. Moreover, this memory deficiency was accompanied by reduced activity not only in the CA1, where the astrocytes are modulated, but also in the ACC, three weeks after manipulation. This temporal association, however, does not necessarily indicate causality, and two possible explanations can be offered: 1) that astrocytic disruption induces a long-term process whose consequences are only observed weeks later, or 2) that it acutely impairs the acquisition of remote (but not recent) memory. We exclude the first option below, and then test the latter.

### Modulation of CA1 astrocytes has no effect on hippocampal neurogenesis

Our findings of intact recent memory followed by impaired remote memory and reduced hippocampal activity could suggest that astrocytic modulation during acquisition initiated a long-term process that took weeks to convey its effect. One example for such a process could be hippocampal neurogenesis occurring between the recent and the remote time points, which had been repeatedly shown to impair remote memory^36–38^. Based on the existence of a sparse projection from dorsal CA1 to DG^39^, and potential indirect influence via the entorhinal cortex, we sought to examine whether astrocytic manipulation induced changes in neurogenesis that can explain the deterioration in memory performance. To tag newborn cells, we administered BrdU (100mg/kg, i.p.), together with the CNO or saline injection, to GFAP::hM4Di mice 30min before FC acquisition, and then another dose 2hr after training. Brains from mice tested for recent contextual memory retrieval were stained for BrdU, tagging the cells added to the DG since the previous day (Figure 2J). No changes in proliferation (Figure 2K) or in the number of cells expressing Doublecortine (DCx), a marker of young neurons 3 days to 3 weeks old (Figure 2L), were observed. Brains collected after remote recall were also stained for BrdU (Figure 2M). No changes in the survival of cells formed on the day of acquisition three weeks previously, or their differentiation fate (determined by co-staining with the neuronal marker NeuN) were observed (Figure 2N). Additionally, no change in the number of young neurons born during these three weeks, marked by DCx, was observed (Figure 2O). CNO application in GFAP::eGFP control mice had no effect on neurogenesis 24 hours or 21 days later (Figure S3L-Q).

To conclude, astrocytic manipulation in CA1 had no effect on hippocampal neurogenesis, and thus an alternative mechanism to the specific impairment of remote memory was subsequently investigated.

### Gi pathway activation in CA1 astrocytes prevents the recruitment of the ACC during memory acquisition

Our findings show that remote memory performance and cFos levels in CA1 and ACC are temporally associated, i.e. when remote recall is low so are cFos levels at the time of recall, but it is challenging to conclude which phenomenon underlies the other. Furthermore, the temporal distance between the appearance of these phenotypes and the time of manipulation three weeks earlier, makes it hard to determine exactly when they were induced. We thus tested the immediate effects of CA1 astrocytic modulation on neuronal activity at the time of memory acquisition. GFAP::hM4Di mice were injected with saline or CNO before FC acquisition, and brains were collected 90 minutes later (Figure 3A). CNO administration had no effect on immediate freezing in response to the foot-shock (Figure S4A). To control for the general effect of astrocytic manipulation on neuronal activity, independent of learning, we manipulated astrocytes not only in fear-conditioned but also in home-caged mice. cFos expression was quantified in 5 brain regions known to be involved in memory: CA1, ACC, BLA, DG and RSC (Figure S4B). Fear conditioning acquisition induced an overall increase in cFos expression in the CA1, ACC and BLA (F_(1,21)_=8.097 p<0.01; F_(1,17)_=5.071 p<0.05; F_(1,16)_=9.067 p<0.01; respectively)(Figure 3B-D; S4C-F), but not in the DG and RSC (Figure S4E,G,H). Astrocytic manipulation in CA1 did not significantly affect local neuronal cFos expression in this region in either home-caged or fear-conditioned mice (Figure 3B,C; S4C).

**Figure 3.**
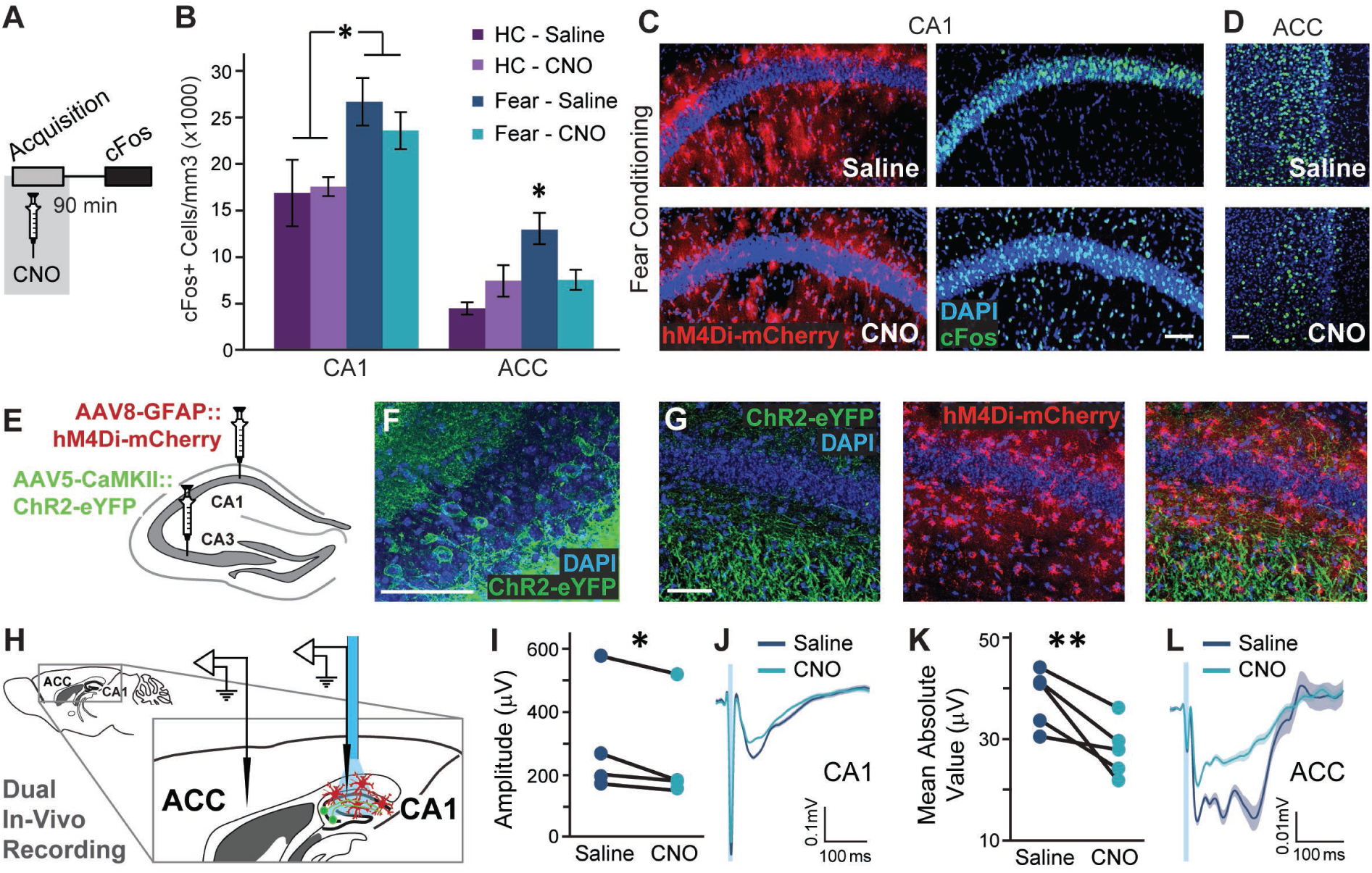
Astrocytic Gi activation in the CA1 prevents the recruitment of the ACC during memory acquisition, and inhibits CA1 to ACC communication. (**A**) GFAP::hM4Di mice were injected with CNO (n=9) or Saline (n=9) 30 minutes before fear conditioning, and brains were removed 90 minutes later for cFos quantification. (**B**) Fear-conditioned GFAP::hM4Di mice showed increased cFos levels in the CA1 compared to home-caged mice (p<0.01), but CNO administration had no effect on either group. cFos levels in the ACC were increased in GFAP::hM4Di that underwent conditioning after being injected with Saline (p<0.05), but not in CNO-injected mice. Data presented as mean ± SEM. Representative images of hM4Di (red) and cFos (green) in the CA1 (**C**) and ACC (**D**) of fear-conditioned mice are presented. cFos-expressing astrocytes are observed below and above the CA1 pyramidal layer in CNO-treated mice. (**E**) AAV5-CaMKII::Channelrhodopsin-2(ChR2)-eYFP was injected into the CA3 and AAV8-GFAP::hM4Di-mCherry into CA1. (**F**) ChR2-eYFP was expressed in the soma of CA3 pyramidal cells. (**G**) The ChR2-expressing axons (green) are observed in the CA1 *stratum radiatum*, and hM4Di-expressing astrocytes (red) are observed in CA1. (**H**) Experimental setup: Light was applied to CA1 in anesthetized mice. The response to Schaffer collaterals optogenetic stimulation was simultaneously recorded in the CA1 and ACC, after Saline administration, followed by CNO administration. (**I**) The response in the CA1 to Schaffer collaterals optogenetic stimulation had a smaller amplitude under Gi-pathway activation by CNO in CA1 astrocytes (n= 4 mice; p<0.05). (**J**) The average responses from one mouse under Saline and then under CNO are presented (average in a bold line, SEM in shadow, blue light illumination in semi-transparent blue). (**K**) A downstream response of CA1 activation by Schaffer collaterals optogenetic stimulation was detected in the ACC. The mean absolute value of the complex ACC response was found to have significantly smaller amplitude under Gi-pathway activation by CNO in CA1 astrocytes (n= 5 mice; p<0.01). (**L**) The average responses from one mouse under Saline and then under CNO are presented (average in a bold line, SEM in shadow). All scale bars=50μm.

Surprisingly, Gi activation on CA1 astrocytes significantly reduced the learning-induced elevation in cFos expression in the ACC, where no direct manipulation took place (Figure 3B,D; S4D). This result is reflected by a significant treatment by behavior interaction (F_(1,17)_=5.036, p<0.05; FC-saline vs. FC-CNO post-hoc, p<0.05). The effect was specific to the ACC, and was not observed in the other non-manipulated regions that were tested, like the BLA, DG or RSC (Figure S4E-H).

The finding that astrocytic Gi pathway activation in CA1 prevented the recruitment of the ACC during learning, suggests a functional CA1→ACC connection, which can be modulated by hippocampal astrocytes. The existence of a monosynaptic CA1→ACC projection had been demonstrated^40^, but to the best of our knowledge a functional connection beyond correlated activity^41^ was never reported. To generate synaptic input to CA1 we expressed channelrhodopsin-2 (ChR2) in CA3 (Figure 3E,F), a major CA1 input source. ChR2-expressing axons from CA3 were observed in the CA1 stratum radiatum, and hM4Di was concomitantly expressed in CA1 astrocytes (Figure 3E,G). Importantly, no fluorescence was detected in the ACC, as there is no direct CA3→ACC projection (Figure S4I,J). Light was applied to CA1 in anesthetized mice, and the fiber was coupled to an electrode recording the neuronal response in CA1 to Schaffer collaterals optogenetic stimulation (Figure 3H). A second electrode was placed in the ACC to record the downstream response to CA1 activation (Figure 3H, Figure S4I,J). Recordings were performed after i.p. saline administration and then after i.p. CNO administration. Optogenetic stimulation of the Schaffer collaterals induced a local response in CA1, which was mildly but significantly reduced by CNO injection (p<0.05, paired t-test; Figure 3I,J). Astrocytic manipulation in the CA1 had a dramatic effect on the downstream response in the ACC to stimulation of the Schaffer collaterals, reflected by a significantly attenuated fEPSP following CNO administration (p<0.01, paired t-test; Figure 3K,L). These results suggest that astrocytic manipulation in CA1 can indeed modulate the functional connectivity from CA1 to ACC.

We show that Gi pathway activation in CA1 astrocytes during fear memory acquisition prevented the recruitment of the ACC, without having a significant effect on local neuronal activity in the CA1, and that CA1 astrocytes can indeed modulate the functional CA1→ACC connectivity. These findings could suggest that astrocytic manipulation selectively blocked the activity of CA1 neurons projecting to the ACC, resulting in a significant effect on ACC activity, but only a mild influence on total CA1 activity.

### Gi activation in CA1 astrocytes during memory acquisition specifically prevents the recruitment of CA1 neurons projecting to ACC

From our findings that Gi activation in CA1 astrocytes during learning prevented the recruitment of the ACC, and that CA1 astrocytes are able to modulate CA1→ACC functional connectivity, we drew the hypothesis that astrocytic Gi activation can selectively prevent the recruitment of CA1 neurons projecting to the ACC, without similarly affecting other CA1 neurons.

To directly test this hypothesis we tagged these projection neurons, measured their recruitment during memory acquisition, and how it is affected by astrocytic Gi activation. Mice were bilaterally injected with a retro-AAV inducing the expression of the Cre recombinase in excitatory neurons (AAV-retro-CaMKIIα::Cre) into the ACC, and an additional Cre-dependent virus inducing the expression of GFP (AAV5-ef1α::DIO-GFP) into CA1 (Figure 4A). AAV8-GFAP::hM4Di-mCherry was simultaneously injected into the CA1, to allow astrocytic manipulation (Figure 4A). Together, these three vectors induced the expression of GFP only in CA1 neurons projecting to the ACC, and of hM4Di in hippocampal astrocytes (Figure 4B-C). These mice were injected with saline or CNO 30 minutes before FC acquisition or in their home cage, and brains were collected 90 minutes later (Figure 4D *top*). As in the previous experiment, CNO administration had no effect on immediate freezing following shock administration (Figure 4D *bottom*), FC acquisition induced an overall increase in cFos expression in the CA1 (F_(1,21)_=12.9 p<0.05), and astrocytic modulation was not sufficient to significantly reduce CA1 cFos expression (Figure 4E). Furthermore, as before, modulation of CA1 astrocytes significantly reduced the learning-induced elevation in ACC cFos expression (Figure 4E; p<0.05, t-test).

**Figure 4.**
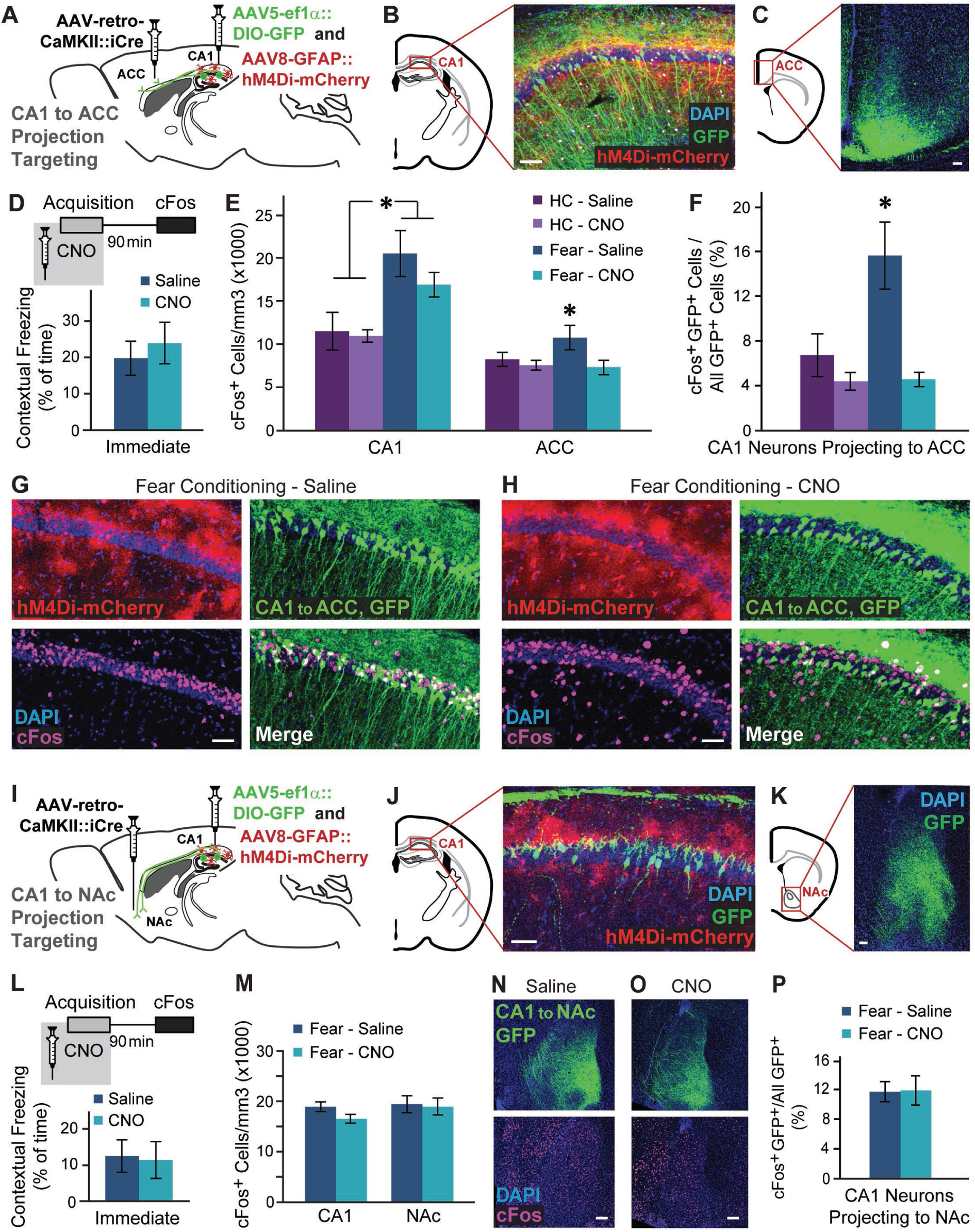
Gi pathway activation in CA1 astrocytes during memory acquisition specifically prevents the recruitment of CA1 neurons projecting to ACC. (**A**) AAV-retro-CaMKII::Cre was injected into the ACC, and AAV5-ef1α::DIO-GFP together with AAV8-GFAP::hM4Di-mCherry were injected into CA1. (**B**) Together, these three vectors induced the expression of GFP (green) in CA1 neurons projecting to the ACC, and hM4Di (red) in CA1 astrocytes. (**C**) GFP-positive axons of CA1 projection neurons are clearly visible in the ACC. (**D**) Mice expressing GFP in ACC-projecting CA1 neurons and hM4Di in their CA1 astrocytes that were injected with CNO (n=8) or Saline (n=7) 30 minutes before FC showed similar immediate freezing following shock administration. (**E**) Fear-conditioned mice showed increased cFos levels in the CA1 compared to home-caged mice (p<0.05), with no effect for CNO administration. cFos levels in the ACC were increased in mice that underwent conditioning after being injected with Saline (p<0.05), but not in CNO-injected mice. (**F**) Fear-conditioned mice injected with Saline showed an >130% increase in the percent of CA1 cells projecting into the ACC that express cFos, compared to home-caged mice (p<0.05). CNO administration completely abolished the recruitment of these cells during learning. Representative images of hM4Di in astrocytes (red), GFP in ACC-projecting CA1 neurons (green) and cFos (pink) in the CA1 of Saline-(**G**) or CNO-(**H**) injected mice are presented. (**I**) AAV-retro-CaMKII::Cre was injected into the NAc, and AAV5-ef1α::DIO-GFP together with AAV8-GFAP::hM4Di-mCherry were injected into CA1. (**J**) Together, these three vectors induced the expression of GFP (green) in CA1 neurons projecting to the NAc, and hM4Di (red) in CA1 astrocytes. (**K**) GFP-positive axons of CA1 projection neurons are clearly visible in the NAc. (**L**) Mice expressing GFP in NAc-projecting CA1 neurons and hM4Di in their CA1 astrocytes that were injected with CNO (n=10) or Saline (n=8) 30 minutes before FC showed similar immediate freezing following shock administration. (**M**) Fear-conditioned mice showed no effect for CNO administration on activity in the CA1, and cFos levels in the NAc were similarly unaltered. Representative images of GFP in the axons of NAc-projecting CA1 neurons (green) and cFos (pink) in the NAc of Saline-(**N**) or CNO-(**O**) injected mice are presented. (**P**) CNO administration had no effect on the activity of CA1 cells projecting into the NAc. All scale bars=50μm. Data presented as mean ± SEM.

When specifically observing the sub-population of ACC-projecting CA1 neurons (CA1→ACC), these cells were found to be dramatically recruited during memory acquisition, and astrocytic modulation significantly reduced the learning-induced cFos elevation in this population (Figure 4F). Specifically, in saline treated mice, more than 15% of the CA1→ACC cells expressed cFos following learning, whereas in CNO-treated GFAP::hM4Di mice less than 5% CA1→ACC cells were active after learning (Figure 4F-H), a level as low as that of home-caged mice (Figure 4F; S5A,B). This effect resulted in a significant treatment by behavior interaction (F_(1,20)_=5.79, p<0.05; FC-saline vs. FC-CNO post-hoc p<0.05).

Finally, to test the specificity of our findings, we then similarly tested an additional monosynaptic projection from the CA1, terminating at the Nucleus Accumbens (NAc). Mice were bilaterally injected with AAV-retro-CaMKII::Cre into the NAc, as well as AAV5-ef1α::DIO-GFP and AAV8-GFAP::hM4Di-mCherry into CA1 (Figure 4I), to tag CA1 neurons projecting to the NAc, and activate the Gi pathway in astrocytes (Figure 4I-K). These mice were injected with saline or CNO before FC acquisition, and brains were collected 90 minutes later (Figure 4L *top*). As in the previous experiment, CNO administration had no effect on immediate freezing (Figure 4L *bottom*), and astrocytic modulation was not sufficient to significantly affect CA1 cFos expression (Figure 4M). Importantly, modulation of CA1 astrocytes had no effect on cFos expression after learning in the NAc (Figure 4M-O). When we specifically tested cFos expression in the sub-population of NAc-projecting CA1 neurons (CA1→NAc), we found that astrocytic modulation had no effect on their activity (Figure 4P, Figure S5C,D).

To conclude, we found that astrocytic inhibition in the CA1 specifically prevented the recruitment of CA1→ACC projecting neurons during memory acquisition. The fact that the inhibition of this projection is induced by the same manipulation that specifically impairs remote memory acquisition, suggests that the activity of CA1→ACC neurons during memory acquisition is necessary for remote recall.

## DISCUSSION

Recent years have seen a burst in discoveries of hitherto unknown elaborate roles for astrocytes in the modulation of neuronal activity and plasticity^42^. In this work, we used advanced tools to modulate CA1 astrocytes, and show for the first time that these cells can confer specific effects on neurons in their vicinity based on the projection target of these neurons. Specifically, astrocytic Gi activation during memory acquisition impairs remote, but not recent, memory retrieval. Another novel finding we present is a massive recruitment of ACC-projecting CA1 neurons during memory acquisition, a process specifically inhibited by astrocytic manipulation, thus preventing a successful recruitment of the ACC during learning.

Previous evidence suggests that astrocytes could have projection-specific effects, based on either the input source or the output target of their neighboring neurons, but with some caveats. For example, in the central amygdala, astrocytic activation depressed inputs from the basolateral amygdala, and enhanced inputs from the central-lateral amygdala^13^. However, since the former projection is excitatory, and the latter inhibitory, this finding could reflect specificity to the secreted neurotransmitter, rather than to the source structure sending these projections. Similarly, two subclasses of astrocytes in the dorsal striatum were shown to specifically modulate either the direct or the indirect pathways^7^. Nonetheless, since the populations of striatal medium spiny neurons from which these two projections originate differ genetically (expressing either the D1 or D2 dopamine receptors), again it is impossible to determine whether the specificity astrocytes show in their effects on these cells stems from their surface protein expression or their projection target. Here, we show for the first time differential effects of astrocytic modulation on CA1 pyramidal cells, based exclusively on their projection target. These cells may differ from other CA1 cells in the configuration of input they receive, their activity pattern, and possibly even in hitherto unidentified genetic properties.

The leading hypothesis in the memory field was that the hippocampus has a time-limited role in memory – required for acquisition and recent recall, and becoming redundant for remote recall, being replaced by frontal cortices^2^. However, this temporal separation between the hippocampus and frontal cortex is not so rigid. For example, we and others have shown that the hippocampus is still critically involved in the consolidation and retrieval of remote memory (e.g.^31, 33, 43–45^). Current research now attempts to define the temporal dynamics in the different brain regions underlying remote memory^33, 45^. The evidence regarding the role of frontal cortices during acquisition is mixed: Inhibition of medial entorhinal cortex input into the PFC during acquisition specifically impaired remote memory^46^. Conversely, chemogenetic inhibition of the PFC during acquisition had no effect on remote recall, nor did optogenetic activation of the PFC neurons that were active at the time of acquisition during remote recall^47^. The role of the ACC in remote memory retrieval was repeatedly demonstrated by the finding that ACC inhibition during recall impairs remote but not recent memory in multiple tasks ^e.g.30–32, 48, 49^, and that sleep deprivation after acquisition, which also impairs only remote memory, reduces ACC recruitment during recall^50^. However, the time-point at which the ACC is recruited to support remote memories was never defined. Here, we show that the ACC is recruited at the time of initial acquisition, but the significance of this early activity is only revealed at the remote recall time point. We further demonstrate, for the first time, massive recruitment of ACC-projecting cells in the CA1 during learning, and show that specific inhibition of this projection at this time-point by astrocytes prevents the engagement of the ACC during acquisition, and results in impaired remote (but not recent) memory. When a non-specific CA1 inhibition is induced by direct neuronal Gi pathway activation, both recent and remote memory is impaired.

We have previously shown that astrocytic activation in CA1 can result in increased neuronal activity in a task-dependent manner and enhance recent memory recall. In this work, we reveal another novel capacity of astrocytes – to affect their neighboring neurons based on their projection target. This finding further expands the repertoire of sophisticated ways by which astrocytes shape neuronal networks and consequently high cognitive function.

## ACKNOWLEDGMENTS

We thank the entire Goshen lab for their support. AK is supported by the ELSC graduate students scholarship. AA is supported by the Adams fellowship. This project has received funding from the European Research Council (ERC) under the European Union’s Horizon 2020 research and innovation programme (grant agreement No 803589 to IG). IG is also supported by the Israel Science Foundation (ISF grant No. 1815/18), the Israeli Centers of Research Excellence Program (center No. 1916/12), and the Canada-Israel grants (CIHR-ISF, grant No. 2591/18). ML is a Sachs Family Lecturer in Brain Science and is supported by the ISF (ISF grant No. 1024/17) and the Einstein Foundation. We thank Yaniv Ziv, Ami Citri, Inna Slutsky, and Adi Doron for the critical reading of the manuscript.

## METHODS

### Subjects

Male C57BL6 mice, 6-7 weeks old (Harlan) were group housed on a 12-hr light/dark cycle with *ad libitum* access to food and water. Experimental protocols were approved by the Hebrew University Animal Care and Use Committee and met guidelines of the National Institutes of Health guide for the Care and Use of Laboratory Animals.

### Virus Production

The pAAV-CaMKII-eGFP plasmid was made by first replacing the CMV promoter in a pAAV-CMV-eGFP vector with the CaMKII promoter. The pAAV-CaMKII-iCre plasmid was made by replacing the eGFP gene in the above plasmid with the coding region of iCre (Addgene 51904). Both pAAV-CaMKII-eGFP and pAAV-CaMKII-iCre plasmids were then packaged into AAV2-retro serotype viral vector. Similarly, pAAV-EF1-DIO-eGFP (Addgene 37084) plasmid was used to make the AAV5-EF1-DIO-eGFP viral vector. The above viral vectors were prepared at the ELSC Vector Core Facility (EVCF) at the Hebrew University of Jerusalem.

### Stereotactic Virus Injection

Mice were anesthetized with isoflurane, and their head placed in a stereotactic apparatus (Kopf Instruments, USA). The skull was exposed and a small craniotomy was performed. To cover the entire dorsal CA1, mice were bilaterally microinjected using the following coordinates: For CA1 (two sites per hemisphere), site 1: anteroposterior (AP), −1.5mm from bregma, mediolateral (ML), ± 1mm, dorsoventral (DV), −1.55mm; site 2: AP −2.5mm, ML ±2mm, DV −1.55mm. For ACC: AP 0.25mm, ML ± 0.4mm, DV −1.8mm. For Schaffer collaterals optogenetic activation, mice were bilaterally microinjected into the CA3 using the following coordinates: AP −1.85, ML +/-2.35, DV −2.25. All microinjections were carried out using a 10µl syringe and a 34 gauge metal needle (WPI, Sarasota, USA). The injection volume and flow rate (0.1μl/min) were controlled by an injection pump (WPI). Following each injection, the needle was left in place for 10 additional minutes to allow for diffusion of the viral vector away from the needle track, and was then slowly withdrawn. The incision was closed using Vetbond tissue adhesive. For postoperative care, mice were subcutaneously injected with Rimadyl (5mg per kg). See supplementary materials for the list of all vectors.

### Immunohistochemistry

3 weeks post-injection mice were transcardially perfused with cold PBS followed by 4% paraformaldehyde (PFA) in PBS. The brains were extracted, postfixed overnight in 4% PFA at 4°C, and cryoprotected in 30% sucrose in PBS. Brains were sectioned to a thickness of 40μm using a sliding freezing microtome (Leica SM 2010R) and preserved in a cryoprotectant solution (25% glycerol and 30% ethylene glycol, in PBS). Free-floating sections were washed in PBS, incubated for 1 h in blocking solution (1% bovine serum albumin (BSA) and 0.3% Triton X-100 in PBS), and incubated overnight at 4°C with primary antibodies (See full list of all antibodies below) in blocking solution. For the cFos staining, slices were incubated with the primary antibody for 5 nights at 4°C. Sections were then washed with PBS and incubated for 2 h at room temperature with secondary antibodies (See supplementary materials) in 1% BSA in PBS. Finally, sections were washed in PBS, incubated with DAPI (1µg/ml), and mounted on slides with mounting medium (Flouromount-G, eBioscience, San-Diego, CA, USA).

For neurogenesis staining, BrdU (Sigma 100mg/kg) was injected intraperitoneally together with the CNO injection, as well as 2 hours after the FC training. 90 minutes after recent or remote recall, brains were removed and slices prepared as described above. Sections were fixated in 50% formamide and 50% SSC for 2 hours in 65°C, then incubated in 2N HCl for 30min at 37°C and neutralized in boric acid for 10min. After PBS washes, sections were blocked in 1% BSA with 0.1% Triton-X for 1 hour at room temperature. Sections were incubated with anti-BrdU for 48h at 4°C. Sections were then washed with PBS and incubated with a secondary antibody for 2 hours at room temperature.

### Confocal Microscopy

Confocal fluorescence images were acquired on an Olympus scanning laser microscope Fluoview FV1000 using 4X and 10X air objectives or 20X and 40X oil immersion objectives. Image analysis was performed using either ImageJ (NIH) or Fluoview Viewer version 4.2 (Olympus).

### Behavioral Testing

The FC apparatus consisted of a conditioning box (18×18×30 cm), with a grid floor wired to a shock generator surrounded by an acoustic chamber (Ugo Basile), and controlled by the EthoVision software (Noldus). Three weeks after injections, mice were placed in the conditioning box for 2min, and then a pure tone (2.9 kHz) was sounded for 20sec, followed by a 2sec foot shock (0.4 mA). This procedure was then repeated, and 30sec after the delivery of the second shock mice were returned to their home cages. FC was assessed by a continuous measurement of freezing (complete immobility), the dominant behavioral fear response. Freezing was automatically measured throughout the testing trial by the EV tracking software. To test contextual FC, mice were placed in the original conditioning box, and freezing was measured for 5min. To test auditory-cued FC, mice were placed in a different context (a cylinder-shaped cage with stripes on the walls and a smooth floor), freezing was measured for 2.5min, and then a 2.9kHz tone was sounded for 2.5min, during which conditioned freezing was measured. Mice were tested for recent memory 24hr after acquisition, and for remote memory 21 or 28 days later. In one experiment, an additional remote memory test was performed 66 days after acquisition.

The non-associative place recognition (NAPR) test was conducted in a round plastic arena, 54 cm in diameter or a square or a trapezoid arena with an identical area size (2290cm^2^). Mice were placed in the center of the arena and allowed to freely explore for 5 min. Habituation to the familiar environment (reduced exploration between first and second exposures) was measured using the EthoVision tracking software.

CNO (Tocris) was dissolved in DMSO and then diluted in 0.9% saline solution to yield a final DMSO concentration of 0.5%. Saline solution for control injections also consisted of 0.5% DMSO. 10mg/kg CNO was intraperitoneally (i.p.) injected 30min before the behavioral assays. In the relevant experiments, BrdU (sigma B5002, 100mg/kg) was injected i.p. together with the CNO/Saline and 2 hours after the behavioral experiment.

The results of behavioral tests were analyzed by a two-way ANOVA followed by LSD post-hoc tests, or by Student’s t test, as applicable.

### In-vivo Electrophysiology and Optogenetics

Simultaneous optical stimulation of the Schaffer Collaterals and electrical recordings in CA1 and ACC were performed as follows: Mice were anesthetized with isoflurane, and an optrode (an extracellular tungsten electrode (1MΩ, ∼125µm) glued to an optical fiber (200µm core diameter, 0.39 NA) with the tip of the electrode protruding ∼400µm beyond the fiber end) was used to record local field potential in Stratum Radiatum and illuminate the Schaffer Collaterals. fEPSP recordings were conducted with the optrode initially placed above the dorsal CA1 (AP −1.6mm; ML 1.1mm; DV −1.1mm) and gradually lowered in 0.1mm increments into the Stratum Radiatum (−1.55mm). The optical fiber was coupled to a 473nm solid-state laser diode (Laserglow Technologies, Toronto, Canada) with ∼10mW of output from the fiber. fEPSP recordings from the ACC were similarly performed using an extracellular tungsten electrode (1MΩ, ∼125µm) placed over the ACC (AP 0.25mm; ML 0.4mm; DV −1.3mm) and gradually lowered in 0.1mm increments to 1.8DV. This electrode was dipped in DiI (1mg/1.5ml in 99% ethanol; Invitrogen) to validate the position of the recording site.

To optogenetically activate the Schaffer collaterals, blue light (473 nm) was unilaterally delivered through the optrode. Photostimulation duration was 10 ms, delivered 72 times for each treatment (saline or CNO) every 5 seconds. Saline and CNO were injected i.p. and recording started 30 minutes after each injection.

Recordings were carried out using a Multiclamp 700B patch-clamp amplifier (Molecular Devices). Signals were low-pass filtered at 5 kHz, digitized and sampled through an AD converter (Molecular Devices) at 10 kHz, and stored for off-line analysis using Matlab (Mathworks Inc.). CA1 responses to Schaffer collaterals stimulation were quantified by calculating the amplitude of the fEPSPs relative to the mean baseline levels, defined as a 200ms time window prior to photostimulation. CA1 activation by Schaffer collaterals stimulation resulted in a complex downstream activity in ACC, lasting approximately 400ms. Because this signal had both positive and negative peaks, to estimate the overall magnitude of the response, we have calculated its mean absolute value over the entire 400ms period, from the beginning of photostimulation in CA1.

### Viral vectors

| Vector | Source | Dilution | Injection Volume |
| --- | --- | --- | --- |
| AAV8-GFAP::hM4D(Gi)-mCherry | UNC vector core | 1:10 in PBS<br>When injected alone | 700nl/site |
| | | 1:10 in other vector when injected with AAV5-EF1α::DIO-GFP | |
| AAV8-GFAP::eGFP | UNC vector core | 1:10 in PBS | 700nl/site |
| AAV5-CaMKIIa::hChr2 (H134R)-eYFP | UNC vector core | --- | 250nl/site |
| AAV5-EF1α::DIO-GFP | EVCf | --- | 500nl/site |
| AAV2-retro-CaMKII-iCre | EVCf | --- | 400nl/site |
| AAV5-CaMKII::hM4Di-mCherry | EVCf | --- | 500nl/site |

### Antibodies

| Primary Antibodies |  |  |  |
| --- | --- | --- | --- |
| Antibody | Source | Catalog # | Dilution |
| Chicken anti-GFAP | Millipore | AB5541 | 1:500 |
| Rabbit anti-NeuN | Cell Signaling Technology | 12943 | 1:400 |
| Rat anti-BrdU | Biorad | OBT0030G | 1:200 |
| Guinea pig anti-DCX | Millipore | AB2253 | 1:1000 |
| Rabbit anti c-fos | Synaptic Systems | 226 003 | 1:10,000 |
| Secondary Antibodies |  |  |  |
| Antibody | Conjugated Fluorophore | Source | Dilution |
| Donkey anti-chicken | Alexa Fluor 488 | Jackson Laboratories | 1:500 |
| Donkey anti-rabbit | Alexa Fluor 488 | Jackson Laboratories | 1:500 |
| Donkey anti-goat | Alexa Fluor 594 | Jackson Laboratories | 1:400 |
| Donkey anti-guinea pig | Cy5 | Jackson Laboratories | 1:400 |
| Donkey anti-rat | Cy5 | Jackson Laboratories | 1:400 |

## SUPPLEMENTARY FIGURES

**Figure S1.**
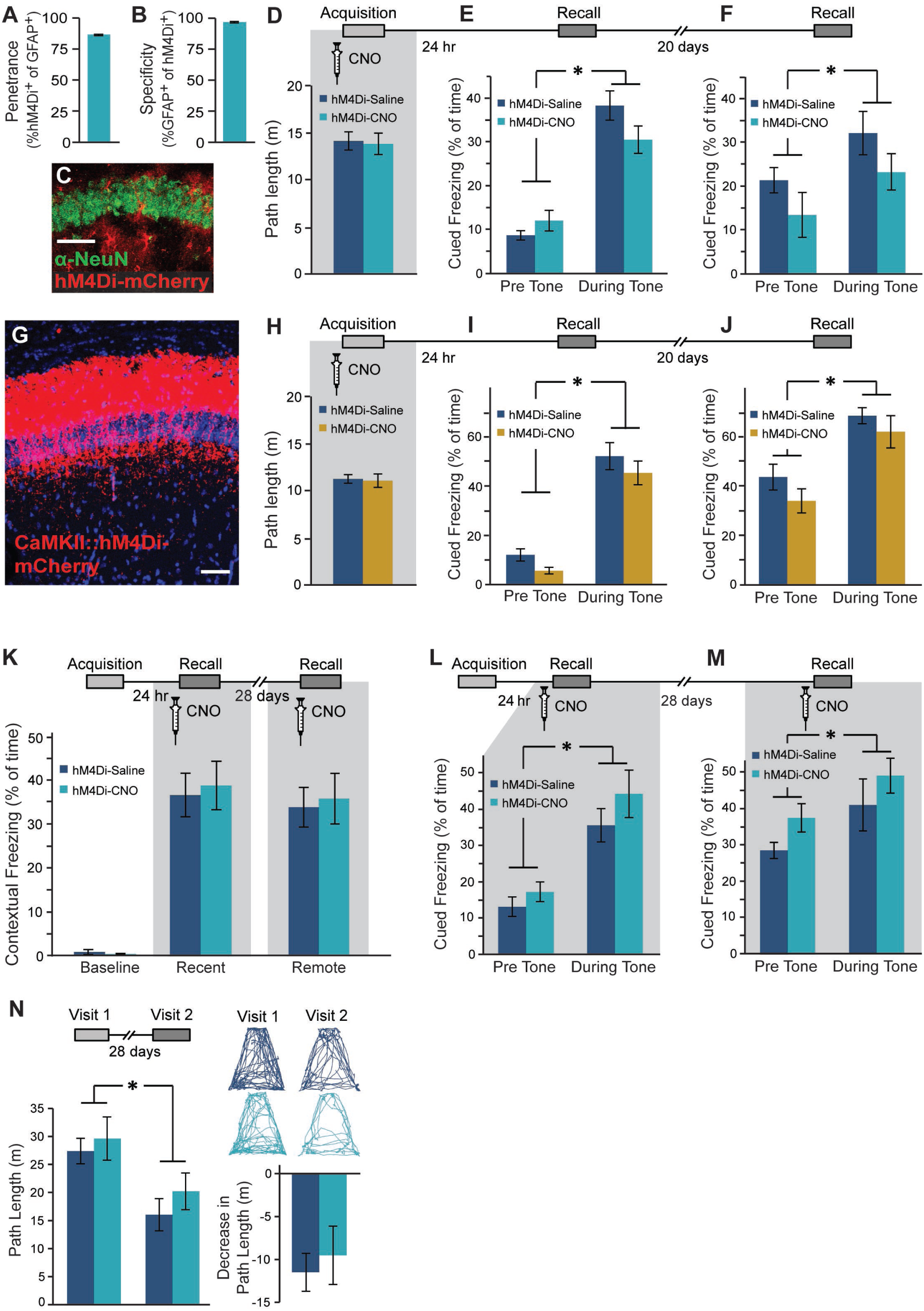
Related to Figure 1. Astrocytic Gi activation in CA1 during learning had no effect on auditory-cued remote memory. Following an injection of AAV8-GFAP::hM4Di-mCherry, hM4Di was expressed in 87% (491/552 cells from 4 mice) of CA1 astrocytes (**A**), with >96% specificity (491/507 cells, from 4 mice)(**B**). (**C**) No co-localization with the neuronal nuclear marker NeuN was detected (scale bar 50µm). GFAP::hM4Di mice were injected with Saline (n=6) or CNO (n=6) 30 min before fear conditioning (FC) acquisition. CNO application before training had no effect on exploration of the conditioning cage (**D**), or on auditory-cued memory recall either 24 hr after acquisition (**E**) or 20 days after that (**F**) in a novel context, with both groups showing increased freezing during tone presentation (p<0.001, p<0.01, respectively). (**G**) Bilateral double injection of AAV5-CaMKIIα::hM4Di-mCherry resulted in hM4Di-mCherry expression in CA1 Neurons only. Scale bar – 50 µm. CaMKIIα::hM4Di mice were injected with either Saline (n=9) or CNO (n=10) 30min before FC acquisition. CNO application before training had no effect on exploration of the conditioning cage (**H**), or on auditory-cued memory recall either 24 hr after acquisition (**I**) or 20 days after that (**J**) in a novel context, with both groups showing increased freezing during tone presentation (p<0.000001, p<0.00001, respectively). (**K**) In a new group of GFAP::hM4Di mice, CNO administration (n=8) only during the recall tests had no effect on either recent or remote memory, compared to Saline-injected controls (n=7). In these mice, CNO administration during recall also had no effect on auditory cued memory either 24 hr after acquisition (**L**) or 20 days after that (**M**), compared to Saline-injected controls. When CNO was not administered during acquisition of the non-associative place recognition task, the GFAP::hM4Di mice (n=6) from Figure 1H showed equivalent performance to controls (n=6; p<0.01)(**N**). Example exploration traces and average Δ are shown (*right*). Data presented as mean ± SEM.

**Figure S2.**
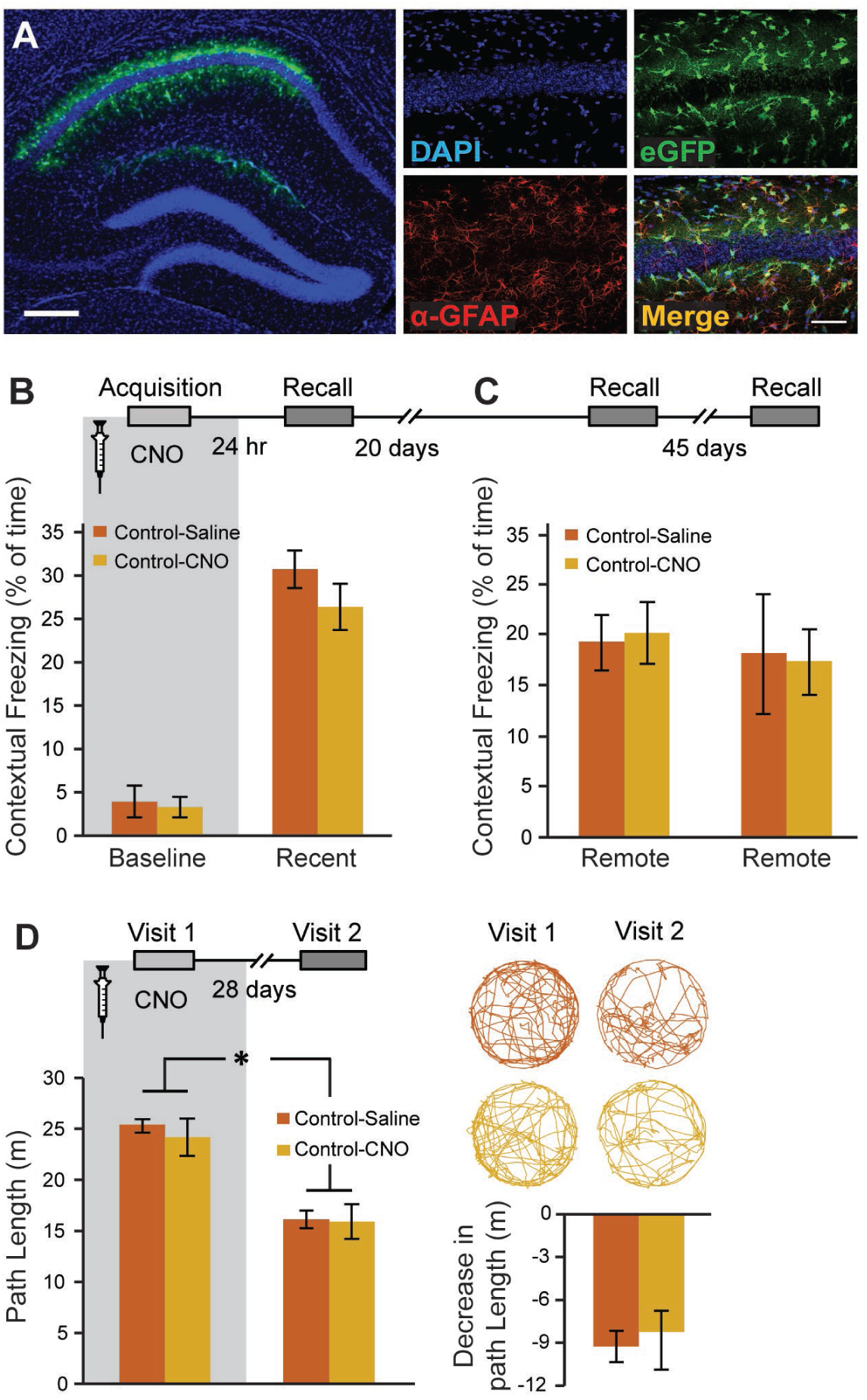
Related to Figure 1. CNO application itself during learning had no effect on remote memory. (**A**) Bilateral double injection of AAV8-GFAP-eGFP resulted in eGFP expression in CA1 astrocytes only. Scale bar – left 300µm, right 50 µm. Mice expressing eGFP in their CA1 astrocytes were injected with either Saline (n=6) or CNO (n=7) 30min before fear conditioning acquisition. CNO administration before training to eGFP-expressing mice had no effect on baseline freezing or recent contextual memory recall one day later (**B**). Neither did CNO have any effect on remote memory 20 days later or 45 days after that (**C**). In the non-associative place recognition test, CNO application before a first visit to a new environment had no effect on remote memory 28 days later (**D**), reflected by a similar decrease (p<0.0001) in the exploration between Saline injected (n=6) and CNO-treated mice (n=7) Example exploration traces and the average change (Δ) following treatment are shown on the right. Data presented as mean ± SEM.

**Figure S3.**
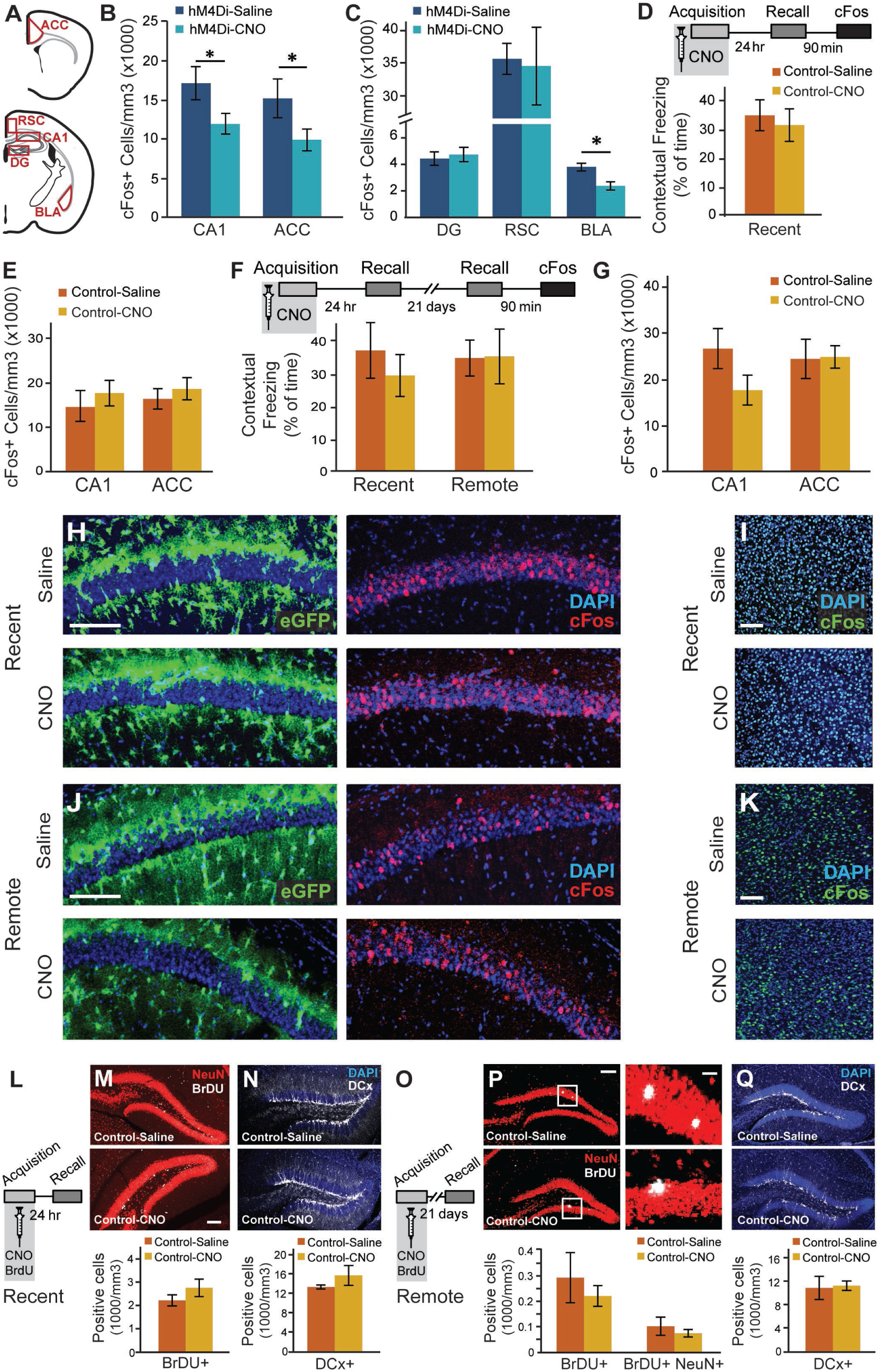
Related to Figure 2. CNO administration during acquisition reduces CA1 and ACC activity at the time of remote recall only in GFAP::hM4Di mice, and does not affect neuronal proliferation, differentiation, or survival. (**A**) Active neurons expressing cFos were quantified in the CA1, ACC, dentate gyrus (DG), retrosplenial cortex (RSC), and basolateral amygdala (BLA). GFAP::hM4Di mice from figure 2A,B that were injected with CNO (n=6) before fear conditioning and showed impaired remote recall compared to Saline controls (n=6), also demonstrated reduced number of cFos expressing neurons in CA1 and ACC (p<0.05 for both)(**B**). No changes in cFos expression in the DG or RSC were observed in these mice, but the reduced fear was accompanied by a significant reduction in cFos expression in the BLA (p<0.05)(**C**). GFAP::eGFP control mice were injected with CNO (n=5) or Saline (n=5) before fear conditioning, and then tested on the next day. No changes were observed in recent memory (**D**) or in the number of neurons active during recent recall in the CA1 or ACC (**E**). Other GFAP::eGFP mice were injected with CNO (n=5) or Saline (n=6) before fear conditioning, and then tested on the next day and again 21 days later. No changes were observed in recent or remote memory (**F**), or in the number of neurons active during remote recall in the CA1 or ACC (**G**). Representative images of GFAP::eGFP (green) and cFos (red in H,J green in I,K) following recent (**H-I**) or remote (**J-K**) recall in the CA1 (**H,J**) and ACC (**I,K**) are presented. (**L**) GFAP::eGFP mice were injected with CNO or Saline together with BrdU before fear conditioning, and then tested on the next day. No changes were observed in stem cell proliferation (Brdu in white)(**M**) or in the number of young, Doublecortine (DCx)-positive neurons (white)(**N**). (**O**) GFAP::eGFP mice were injected with CNO or Saline and BrdU before fear conditioning, and then tested 21 days later. No changes were observed in stem cell proliferation and differentiation (**P**) or in the number of young, DCx-positive neurons (**Q**). Scale bars = 100μm for CA1, ACC and whole DG, 10 μm for zoomed-in cells. Data presented as mean ± SEM.

**Figure S4.**
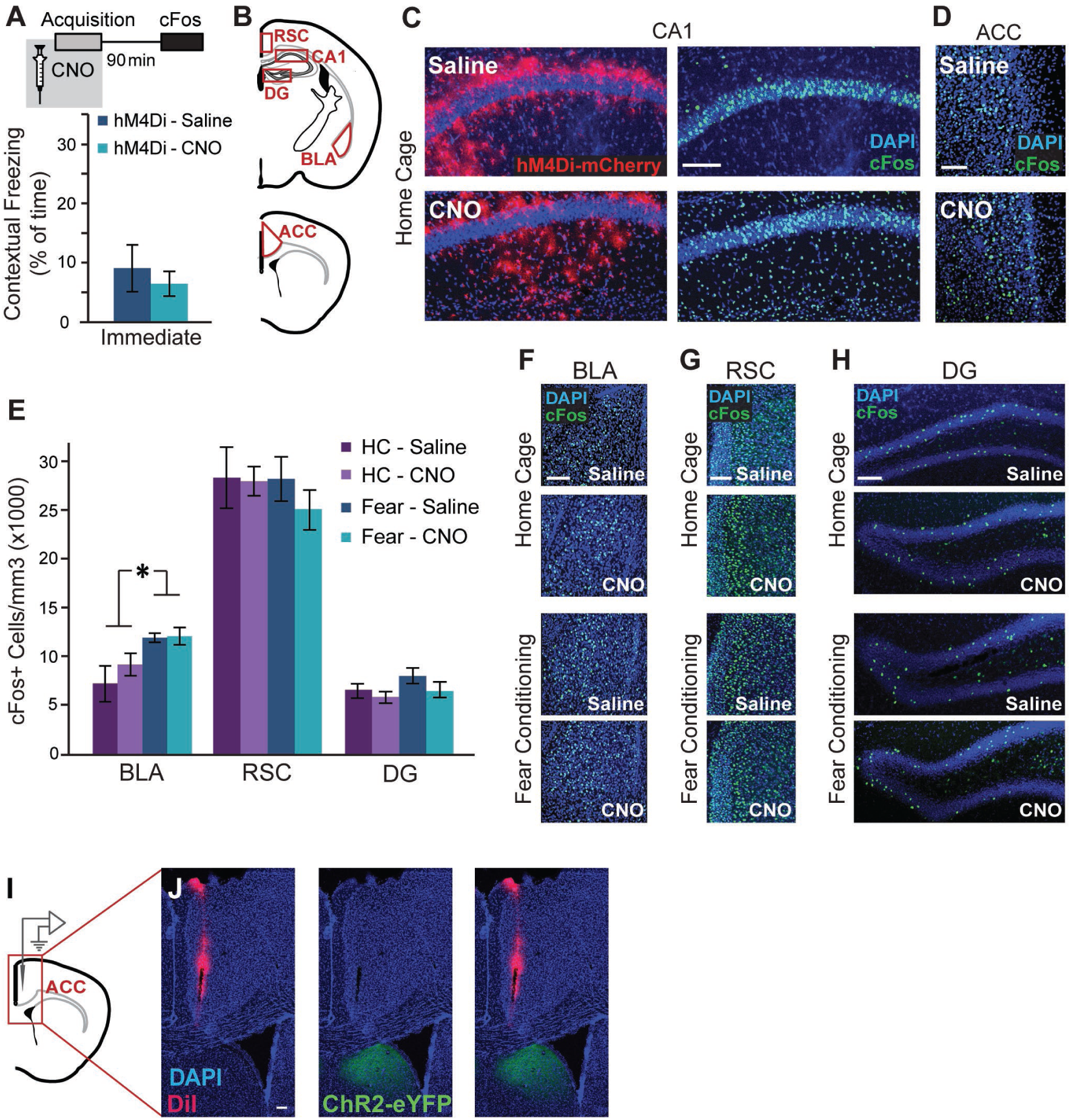
Related to Figure 3. Astrocytic inhibition in CA1 during memory acquisition does not affect the recruitment of the RSC and DG. (**A**) GFAP::hM4Di mice that were injected with CNO (n=9) or Saline (n=9) 30 minutes before fear conditioning showed similar immediate freezing following shock administration to Saline-injected controls. (**B**) Active neurons expressing cFos were quantified in the in the CA1, basolateral amygdala (BLA), ACC, retrosplenial cortex (RSC) and dentate gyrus (DG) of GFAP::hM4Di mice that were injected with CNO (n=9) or Saline (n=9) 30 minutes before fear conditioning, or in home-caged mice (CNO n=4, Saline n=4). (**C**) Representative images of hM4Di (red) and cFos (green) in the CA1 (**C**) and ACC (**D**) of home caged GFAP::hM4Di mice showing no effect of CNO administration on cFos levels. cFos-expressing astrocytes are observed below and above the CA1 pyramidal layer. Scale bars=100μm. (**E**) Fear-conditioned GFAP::hM4Di mice showed increased cFos levels in the BLA compared to home-caged mice (p<0.01), but CNO administration had no effect on either group. Fear-conditioning and CNO administration had no effect on cFos levels in the RSC and DG. Representative images of hM4Di (red) and cFos (green) in the BLA (**F**), RSC (**G**) and DG (**H**) are presented. (**I**) An electrode dipped in DiI was placed in the ACC to record the response to CA1 activation. (**J**) The location of the electrode in the ACC is shown in crimson, and no ChR2-eYFP positive axons (green) are observed in this region. All scale bars = 100μm. Data presented as mean ± SEM.

**Figure S5.**
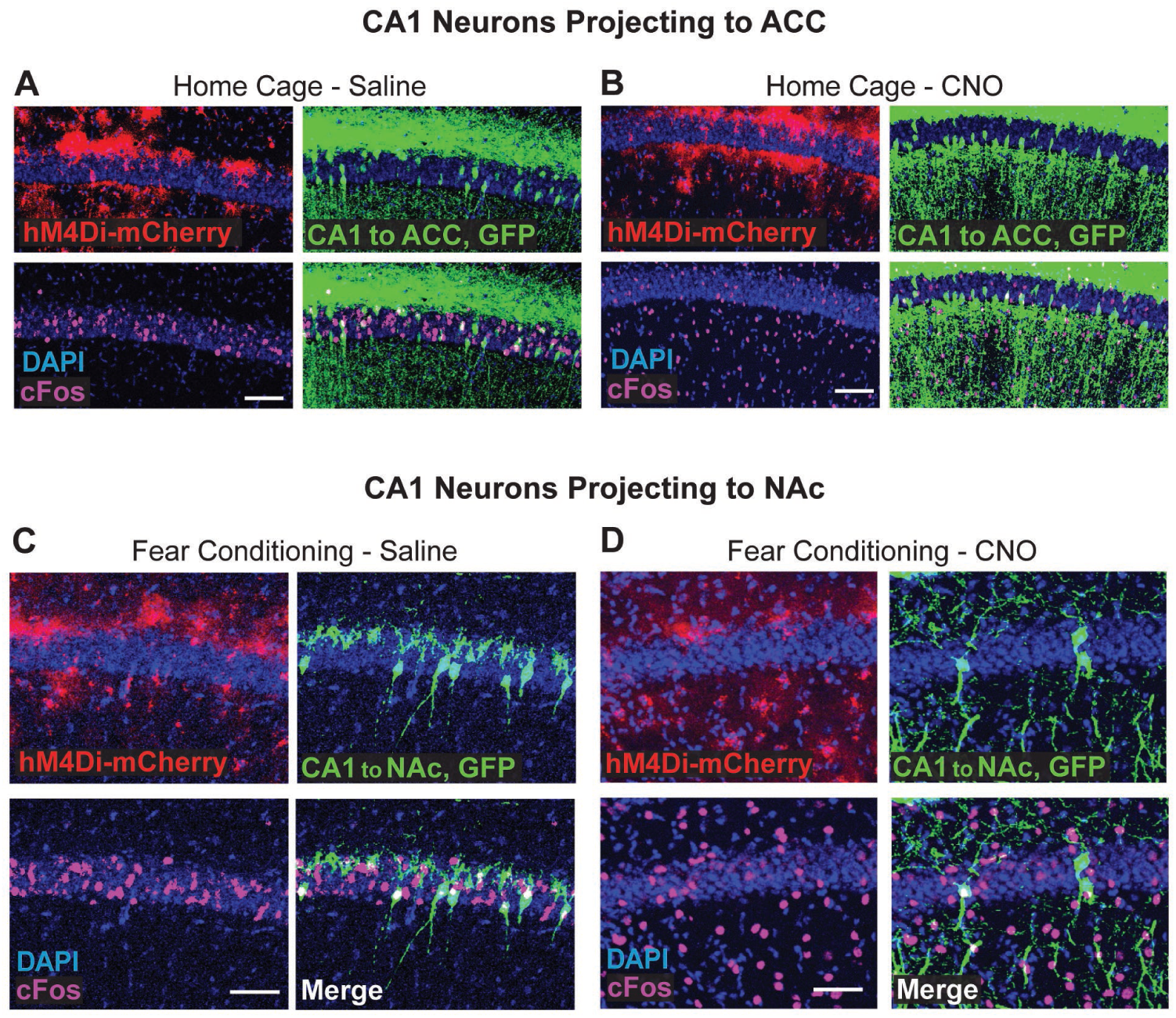
Related to Figure 4. Gi pathway activation in CA1 astrocytes has no effect on cFos expression in home-caged mice. (**A-B**) Representative images of hM4Di in astrocytes (red), GFP in ACC-projecting CA1 neurons (green) and cFos (pink) in the CA1 of Saline- (**A**) or CNO- (**B**) injected home-caged mice are presented. No effect of CNO on cFos levels was observed. (**C-D**) Representative images of hM4Di in astrocytes (red), GFP in NAc-projecting CA1 neurons (green) and cFos (pink) in the CA1 of Saline-(**C**) or CNO-(**D**) injected fear-conditioned mice are presented, showing no effect of the astrocytic manipulation on CA1→NAc neurons activity. All scale bars=50μm.

